# Feedback linking cell envelope stiffness, curvature, and synthesis enables robust rod-shaped bacterial growth

**DOI:** 10.1101/2022.04.01.486519

**Authors:** Salem al-Mosleh, Ajay Gopinathan, Christian Santangelo, Kerwyn Casey Huang, Enrique Rojas

**Affiliations:** John A. Paulson School of Engineering and Applied Sciences, Harvard University, Cambridge, MA 02138, USA; Department of Physics, and SF-CREST: Center for Cellular and Biomolecular Machines, University of California Merced, Merced, CA 95343, USA; Department of Physics, Syracuse University, Syracuse, NY 13210, USA; Department of Bioengineering, and Department of Microbiology and Immunology, Stanford University, Stanford, CA 94305, USA; Chan Zuckerberg Biohub, San Francisco, CA 94158, USA; Department of Biology, New York University, New York, NY, 10003, USA

**Keywords:** cell mechanics, hyperosmotic shock, Gram-negative bacteria, cell envelope, stored growth, envelope softening, adaptation

## Abstract

Bacterial growth is remarkably robust to environmental fluctuations, yet the mechanisms of growth-rate homeostasis are poorly understood. Here, we combine theory and experiment to infer mechanisms by which *Escherichia coli* adapts its growth rate in response to changes in osmolarity, a fundamental physicochemical property of the environment. The central tenet of our theoretical model is that cell-envelope expansion is only sensitive to local information such as enzyme concentrations, cell-envelope curvature, and mechanical strain in the envelope. We constrained this model with quantitative measurements of the dynamics of *E. coli* elongation rate and cell width after hyperosmotic shock. Our analysis demonstrated that adaptive cell-envelope softening is a key process underlying growth-rate homeostasis. Furthermore, our model correctly predicted that softening does not occur above a critical hyperosmotic shock magnitude and precisely recapitulated the elongation-rate dynamics in response to shocks with magnitude larger than this threshold. Finally, we found that to coordinately achieve growth-rate and cell-width homeostasis, cells employ direct feedback between cell-envelope curvature and envelope expansion. In sum, our analysis points to new cellular mechanisms of bacterial growth-rate homeostasis and provides a practical theoretical framework for understanding this process.

**Significance Statement:** The bacterial cell envelope is the critical structure that defines cell size and shape, and its expansion therefore defines cell growth. Although size, shape, and growth rate are important cellular variables that are robust to environmental fluctuations, the feedback mechanisms by which these variables influence cell-envelope expansion are unknown. Here, we explore how *E. coli* cells achieve growth-rate and cell-width homeostasis during fluctuations in osmolarity, a key environmental property. A biophysical model in which the cell envelope softens after an osmotic shock and envelope expansion depends directly on local curvature quantitatively recapitulated all experimental observations. Our study elucidates new mechanisms of bacterial cell morphogenesis and highlights the deep interplay between global cellular variables and the mechanisms of cell-envelope expansion.

---

For bacterial cells, cell growth and cellular morphogenesis are intimately related. For example, to achieve its characteristic rod-shape, *Escherichia coli* must maintain its cell width (Fig. 1) while growing in cell length. Despite its complexity, this process is remarkably robust: bacterial cells maintain cell shape and cell growth with precision across environmental conditions and during dramatic environmental perturbations (1–5). Although much is known about the molecular pathways that are required for increases cell volume and surface area (6, 7), little is known about the homeostatic mechanisms that couple these processes to guarantee stable cell growth and morphogenesis. In particular, it is unknown which global (cell-scale) morphogenetic variables — such as cell width, cell length, or cell growth rate — feedback directly onto the mechanisms that orchestrate cell growth.

**Fig. 1.**
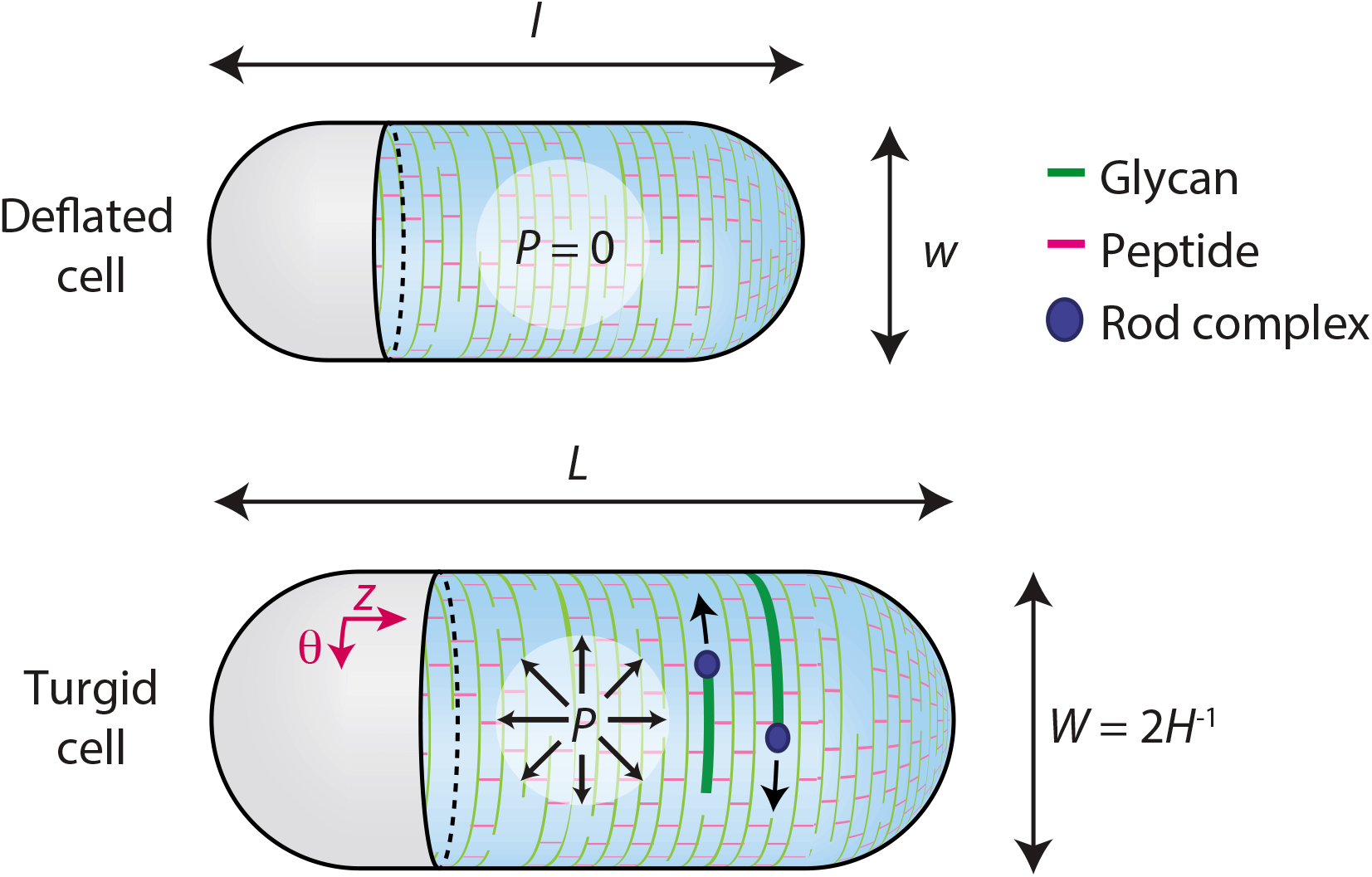
Gram-negative bacterial cell geometry and mechanics. Schematic of a Gram-negative bacterial cell illustrating its rest length *l* and rest width w in the absence of any applied forces such as turgor pressure. During steady-state growth with rate *λ*_0_, turgor pressure *P*(*t*) = *P*_0_ stretches the cell to length *L*(*t*) = *L*(0) exp (*λ*_0_*t*) and width *W*(*t*) = 2*H*^-1^ = *W*_0_, where *H* is the curvature. Synthesis of the peptidoglycan cell wall takes place through the action of Rod complexes (blue circle); glycan strands (green) are inserted in an approximately circumferential direction and crosslinked to old material through short peptides (red).

The cell envelope is the key structure that defines bacterial cell size and shape and therefore the expansion of this surface defines cell growth and morphogenesis. The cell envelope is complex and diverse. In Gram-negative bacteria such as *E. coli,* the envelope is a multi-layered structure that includes the peptidoglycan cell wall and the outer membrane (Fig. 1). Both of these layers contribute to envelope mechanical integrity, while the cell wall is the key structural determinant of cell shape (8).

Synthesis of the cell wall and the outer membrane are carried by sophisticated molecular machineries. Assembly of the cell wall is executed by protein complexes called “Rod complexes” (9) that are distributed throughout the cylindrical (i.e., non-polar) regions of the plasma membrane (Fig. 1) (10). Rod complexes synthesize nascent glycan polymers processively from one end and cross-link them into the existing cell wall via their free peptide stems. Processive synthesis is oriented approximately parallel to the circumference of the cell such that glycans are preferentially oriented circumferentially and peptides are oriented parallel to the cell axis (Fig. 1), resulting in structural anisotropy of the cell wall at the molecular scale. This structural anisotropy is thought to be critical for rod-shape maintenance by stiffening the cell envelope in the circumferential direction (11).

A few paradigmatic cases demonstrate how global morphogenetic variables can regulate molecular-scale cell envelope homeostasis. For example, the positioning of the cell division septum is governed by the spatiotemporal dynamics of the Min proteins, which depend directly on global cell-envelope geometry (12–16). Second, when *E. coli* cells are temporarily constrained to adopt a bent shape within a curved channel and then released, new envelope synthesis by Rod complexes preferentially localized to the inner curvature, which promotes cell straightening (17, 18). It is likely that this localization preference represents feedback from global cell shape via the local geometry of the cell envelope. Similarly, circumferential glycan synthesis is thought to rely on the ability of some components of the Rod complexes to sense membrane curvature (10, 19–22). According to one model, short polymers of the actin homolog MreB, which scaffold other components of the Rod complexes, align circumferentially along the direction of maximal membrane curvature and act as molecular rudders that steer glycan synthesis in this direction (23, 24). These mechanisms likely represent the tip of the iceberg with respect to how global variables feedback onto the molecular-scale machinery of cellular morphogenesis.

Ultimately, cellular morphogenesis is a physical process of cell enlargement and shape changes, yet the ways in which mechanical forces interact with the molecular mechanisms of morphogenesis are not understood. A key question is the role of turgor pressure, another global variable, in promoting cell envelope expansion during cell growth (Fig. 1) (25). In principle, turgor pressure could feed back onto cell envelope expansion in two ways: i) by activating the enzymes that carry out synthesis and/or hydrolysis of the cell envelope, or ii) by direct plastic deformation of the envelope. Moreover, these connections could be mediated through the degree to which the envelope is stretched (the “mechanical strain,” a local variable) or via changes in envelope curvature caused by inflation of the cell by pressure.

To address the role of turgor pressure in cell envelope expansion, in a previous study we measured the growth-rate dynamics of *E. coli* in response to osmotic shocks (acute changes in extracellular osmolarity that alter turgor pressure) (3). The responses were complex: while modest hyperosmotic shocks (<300 mM in magnitude) clearly reduced growth rate, when turgor pressure was re-established by reversing the shock, cells rapidly elongated to the size that they would have had in the absence of the shock (3). That is, the cells appeared to “store growth” during the period of reduced growth rate. Although similar phenomena have been observed in the alga Chara corallina (26), the mechanisms that enable stored growth and the conditions under which it can be achieved are not well understood.

To understand stored growth, and the interplay of cell morphology, cell envelope synthesis, and turgor pressure more broadly, we developed a generic theory for the expansion of a thin pressurized shell (27) that unifies these processes. We validated this model using an extensive data set concerning cellular elongation rate and cell width dynamics after osmotic shocks. The central underlying tenet of our model is “locality”: that the rate of cell envelope expansion depends only on local information including the rate of cell-envelope synthesis, cell-envelope curvature, and mechanical strain in the envelope. We found that the model can only explain stored growth with the inclusion of adaptive cell-envelope softening, which also accelerates recovery of cell width. Moreover, the model successfully predicts a threshold shock magnitude above which stored growth cannot occur due to an instability generated by envelope softening. Beyond this shock magnitude, the model also quantitatively predicts the slope of the experimentally observed linear decrease in elongation rate as a function of shock magnitude. Finally, we found that direct feedback between envelope curvature and envelope expansion is required to explain the transient dynamics of cell length and width after hyperosmotic shock. These results highlight the sophisticated nature of the feedback system that governs cellular morphogenesis, which coordinately controls cell elongation rate and cell width by coupling them to envelope stiffness and synthesis; the specific organization of this feedback system promotes robust and rapid homeostasis of each of these variables.

## Results

### A physical model of surface expansion based on the principle of locality can be adapted to bacterial growth

Our strategy for interrogating bacterial morphogenesis was to derive a mathematical model that is generic enough to capture the rich phenomenology of cell growth dynamics upon perturbation, constrain this model with a broad set of experimental data, and infer principles of morphogenesis from these constraints. Motivated by a large body of experimental support, we chose to derive a coarse-grained model of cell-envelope expansion based on the principle of locality. Here, locality implies that envelope expansion at a given point is only dependent on information within its immediate microscopic vicinity. Such information could be chemical (e.g., local enzyme concentration), mechanical (e.g., mechanical strain in the cell envelope), or geometrical (e.g., curvature of the cell envelope). Within this framework, global (cell-scale) information can only be sensed insofar as it is coupled to local information. For example, the width of a rod-shaped cell could only be sensed indirectly through local curvature since these two variables are inversely related (Fig. 1). Similarly, turgor pressure could be sensed indirectly through local mechanical strain.

Based on this principle, we adapted a differential geometry-based theory for the expansion of thin elastic surfaces to rod-shaped (cylindrical) cell-envelope expansion (28) (Methods, SI). The dynamics of cell growth are described by those of the rest metric tensor, *g_ij_*(*t*); along with the rest curvature tensor, *b_ij_*(*t*), the rest metric tensor specifies the geometry of the cell envelope when there are no external forces applied to it (Fig. S1). In the presence of turgor pressure, the envelope stretches (Fig. 1) and its geometry is described by the metric tensor *G_ij_*(*t*) and curvature tensor *B_ij_*(*t*). Generically, the dynamics of the rest metric tensor may depend on envelope curvature and mechanical strain in the envelope (Fig. S1). We performed a Taylor expansion of metric tensor dynamics to leading orders of curvature and strain, motivated by the fact that the curvature of the cell envelope (inverse radius of curvature) is small compared with the inverse of the size of molecular components such as MreB. We identified four key terms that could affect cell morphology dynamics after hyperosmotic shock (SI):

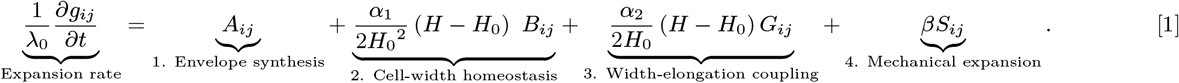

The left-hand side of Eq. 1 is the time derivative of the rest metric tensor, which defines the expansion rates of the cell envelope in both principal directions (longitudinal and circumferential, denoted *z* and *θ*, respectively, in Fig. 1); *λ*_0_ defines the steady-state elongation rate.

Term 1 on the right-hand side of Eq. 1 defines the steady-state anisotropic expansion of the cell surface along the principal curvature directions (SI). Biologically, this term corresponds to the expansion of the cell surface that results from curvature-dependent anisotropic synthesis of envelope material.

Terms 2 and 3 encompass direct feedback between cell envelope curvature, H, and cell surface expansion. H is positive by convention. *H*_0_ = 2/*W*_0_ is the steady-state envelope curvature where *W*_0_ is the steady-state cell width. *α*_1_ and *α*_2_ are proportionality constants. For a cylindrical cell, only the circumferential component of the curvature tensor *B_ij_* is non-zero. Term 2 by itself ensures cell width homeostasis by providing negative feedback between circumferential curvature and circumferential expansion. Term 3 modulates isotropic surface expansion in an envelope curvature-dependent manner. Biologically, these two terms could rely on a single process, for example, curvature-dependent localization of Rod complexes (10).

Term 4 corresponds to irreversible cell-surface expansion driven directly by mechanical strain in the cell envelope, which results from turgor pressure. This term yields anisotropic expansion that is directly proportional to the anisotropy in the strain tensor, *S_ij_* = (*G_ij_* – *g_ij_*)/2, where *β* is a proportionality constant. Notably, this term is not the only mechanism by which envelope expansion could depend on strain: since Terms 1-3 prescribe changes in the rest metric tensor that explicitly depend on envelope shape, and since shape changes in response to mechanical forces, the elongation rate implicitly depends on strain even when *β* = 0.

A minimal version of the model with Terms 1 and 2 alone can achieve stable rod-shaped elongation: a cylinder with constant width equal to 2/*H*_0_ and exponentially increasing length *L*(*t*) = *L*(0) exp (*λ*_0_*t*) is a steady-state solution of Eq. 1 (SI). To determine whether contributions from Terms 3 and 4 are required to explain the dynamics of cell morphogenesis in general, we sought to perturb cell growth from its steady-state behavior. We did so by subjecting cells to hyperosmotic shock, which rapidly changes the curvature and mechanical strain in the cell envelope. This strategy allowed us to directly test whether cell elongation and cell width are coupled due to curvature-dependent cell-envelope expansion, and whether mechanical strain directly drives cell-envelope expansion.

### Stored growth implicates dynamic adaptation of cell envelope stiffness

When *E. coli* cells are perfused constantly with rich medium, they elongate at a steady-state rate of *λ*_0_ = *d*(ln *L*)/*dt* ≈ 0.03 min^-1^. Stored growth occurs when cells are subjected to modest (≤ 300 mM), transient hyperosmotic shocks (5). During hyperosmotic shock, turgor pressure decreases and the cell envelope contracts (Fig. 2A,B), with envelope length governed by linear elasticity:

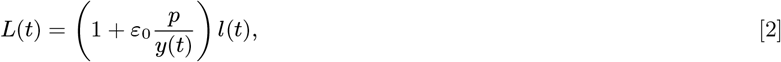

where *l*(*t*) is the rest length of the cell, *p* = *P/P*_0_ is the turgor pressure normalized by its steady-state value, *y*(*t*) = *Y*(*t*)/*Y*_0_ is Young’s modulus of the cell envelope normalized by its steady-state value, and *ε*_0_ is the mechanical strain in the cell envelope prior to the decrease in turgor pressure. In previous work, we measured *ε*_0_ ≈ 0.1 ± 0.03 (standard deviation) (29). Immediately after a hyperosmotic shock, cells elongate slower than before. However, the cells store growth: when the shock is reversed, cells rapidly expand elastically to the size that they would have attained during this period had they never been subjected to the shock (Fig. 2B) (3).

**Fig. 2.**
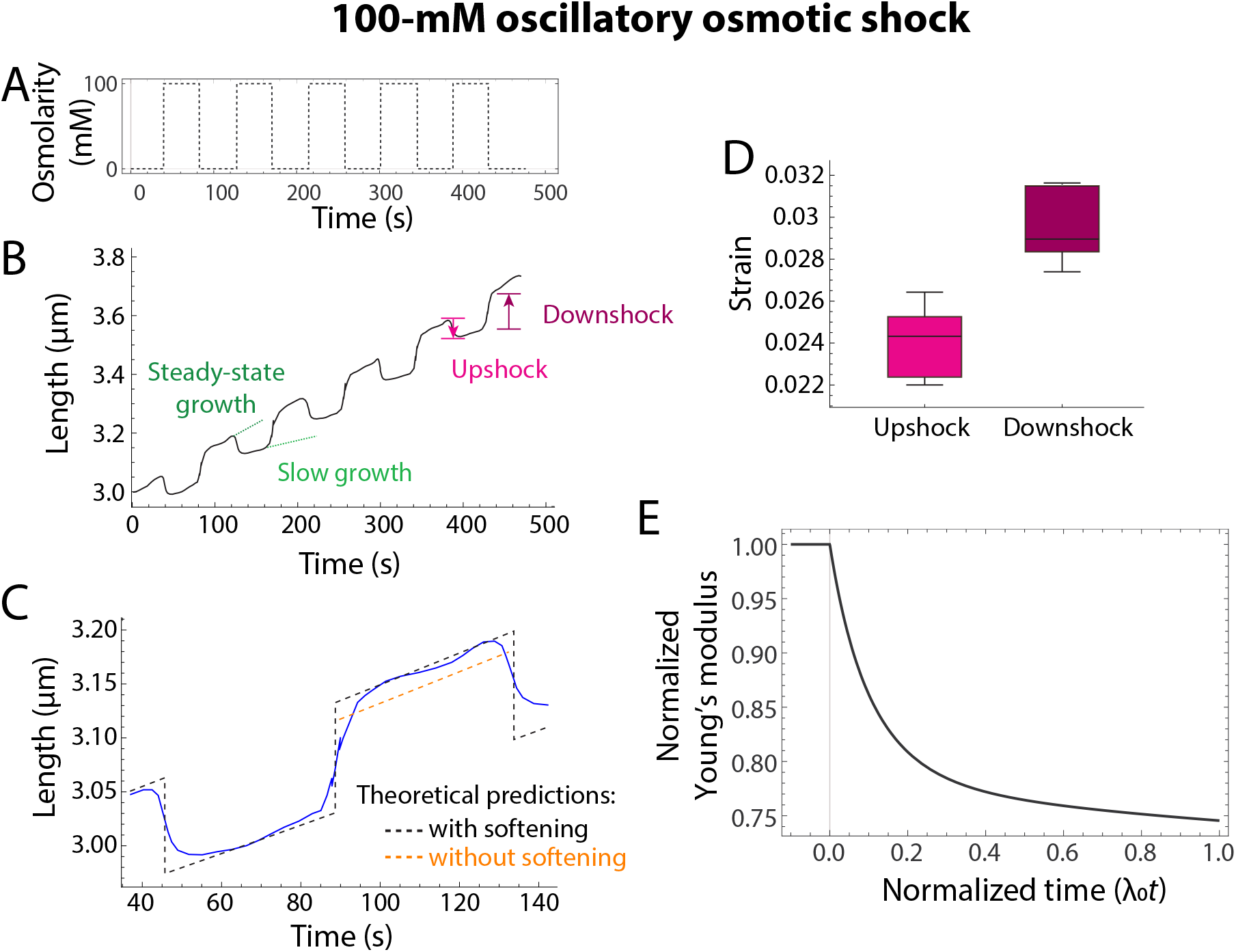
Stored growth during cycles of hyper- and hypoosmotic shocks can be explained by cell envelope softening. (A) Osmolarity as a function of time. (B) Experimentally measured cell length dynamics (black) in response to cycles of 100-mM hyper- and hypoosmotic shocks demonstrate stored growth: immediately after the hyperosmotic shocks, growth rate decreases, yet after the hypoosmotic shocks cell length ends up matching the predicted elongation in unperturbed conditions extrapolated from the initial cell length and growth rate. (C) A closer look at a portion from panel (B). Orange dashed curve shows the predicted dynamics after the hypoosmotic shock in the absence of envelope softening. (D) The mechanical strain induced by the hyperosmotic shocks in (B) is smaller than the strain induced by the hypoosmotic shocks, implying softening of the cell envelope. (E) Model prediction of the envelope softening necessary for stored growth after a 100-mM hyperosmotic shock is in reasonable agreement with the softening inferred from the mechanical strain data as discussed in the text.

To achieve stored growth, the elastic expansion when the shock is reversed (i.e., during the “downshock”, Fig. 2A,B) must necessarily be much larger than the elastic contraction when the shock is applied (“upshock”, Fig. 2A,B) to make up for the slower elongation during the period of reduced turgor pressure (Fig. 2B-D). This difference means that the cell envelope is less stiff at the end of the period of low turgor than directly after the hyperosmotic shock. The condition for stored growth – that the reversal of the shock fully compensates for the length lost during low turgor – leads to an expression for the dynamics of the longitudinal Young’s modulus after hyperosmotic shock, *y*(*t*):

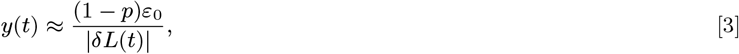

where 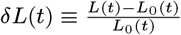 is the relative difference between the cell length, *L*(*t*), and the length that the cell envelope would have had in the absence of perturbation, *L*_0_(*t*). As this difference increases, the longitudinal Young’s modulus decreases. There are no fitting parameters in Eq. 3, hence *y*(*t*) is a direct consequence of the dynamics of *L*(*t*).

The theoretical rate and extent of softening upon hyperosmotic shock can be estimated by solving Eq. 1-3. By substituting the predicted length dynamics into Eq. 3, we can predict the time evolution of longitudinal Young’s modulus after the hyperosmotic shock (Fig. 2E). By comparing the mechanical strains induced by hyperosmotic shock and reversal of the shock (Fig. 2A-D, S2), we find that during the 60 s after a 100-mM hyperosmotic shock, the elastic modulus is reduced by 19 ± 6%, in reasonable agreement with the prediction of 10% after a normalized time *λ*_0*t*_ = 0.033 from our model (Fig. 2E). Remarkably, our model predicts that longitudinal Young’s modulus would decrease by >25% during one cell cycle in the absence of osmoregulation (Fig. 2E).

Envelope softening can be viewed as a mechanical mechanism of sensing global cellular dimensions. Specifically, envelope softening intrinsically records the deviation between global cell length and the length determined by cell-envelope synthesis (i.e., *δL*). Furthermore, softening intrinsically provides a mechanical mechanism to reduce this deviation.

### Cell envelope softening and curvature feedback enable simultaneous homeostasis of cellular elongation rate and cell width

We next hypothesized that cell-envelope softening would affect elongation-rate dynamics on longer time scales after single hyperosmotic shocks (without reversing the shock). To test this hypothesis, we used microfluidics to subject *E. coli* cells growing at steady state to hyperosmotic shocks across a wide range of magnitudes and measured the dynamic responses of cell length over time. We then systematically tested these data against our model (Eq. 1), with and without envelope softening (Eq. 3). Throughout our entire analysis we used a single set of model parameters to explain experimental data (Methods).

By quantifying the relative length difference, *δL*(*t*), and the relative difference between the observed elongation rate and the elongation rate before the shock, *δλ*(*t*) = (λ(*t*) – λ_0_)/λ_0_, we found that the shock caused a short, transient period of reduced elongation rate (Fig. 3A-C), followed by rapid re-establishment of the pre-shock elongation rate (Fig. 3C). In the absence of envelope softening, both the minimal version of our model (the “Minimal model”; Eq. 1, Terms 1 and 2) and the version that includes feedback from envelope curvature (the “Curvature-feedback model”; Eq. 1, Terms 1-3) poorly predicted the experimental cell length dynamics after hyperosmotic shock (Fig. 3A,B). Each model generically predicted that the relative length change, *δL*(*t*), does not approach a constant value (Fig. 3B) and equivalently that the relative elongation rate change, *δλ*(*t*), does not approach zero (Fig. 3C), in contrast to our experimental observations (Fig. 3B,C). When combined with softening (Eq. 3), however, the curvature feedback model accurately captured the length dynamics after both a 400-mM (Fig. 3A-C) and a 200-mM shock (Fig. S3), while the minimal model did not (Fig. 3A-C, S3). The minimal model predicted a smaller initial decrease in elongation rate than in experiments, and the relative rate change was approximately constant over the interval during which the curvature model exhibited adaptation (Fig. 3C). In sum, these findings implicate direct dependence of cell elongation on envelope curvature and point to the critical role of envelope softening during adaptation to changes in turgor pressure.

**Fig. 3.**
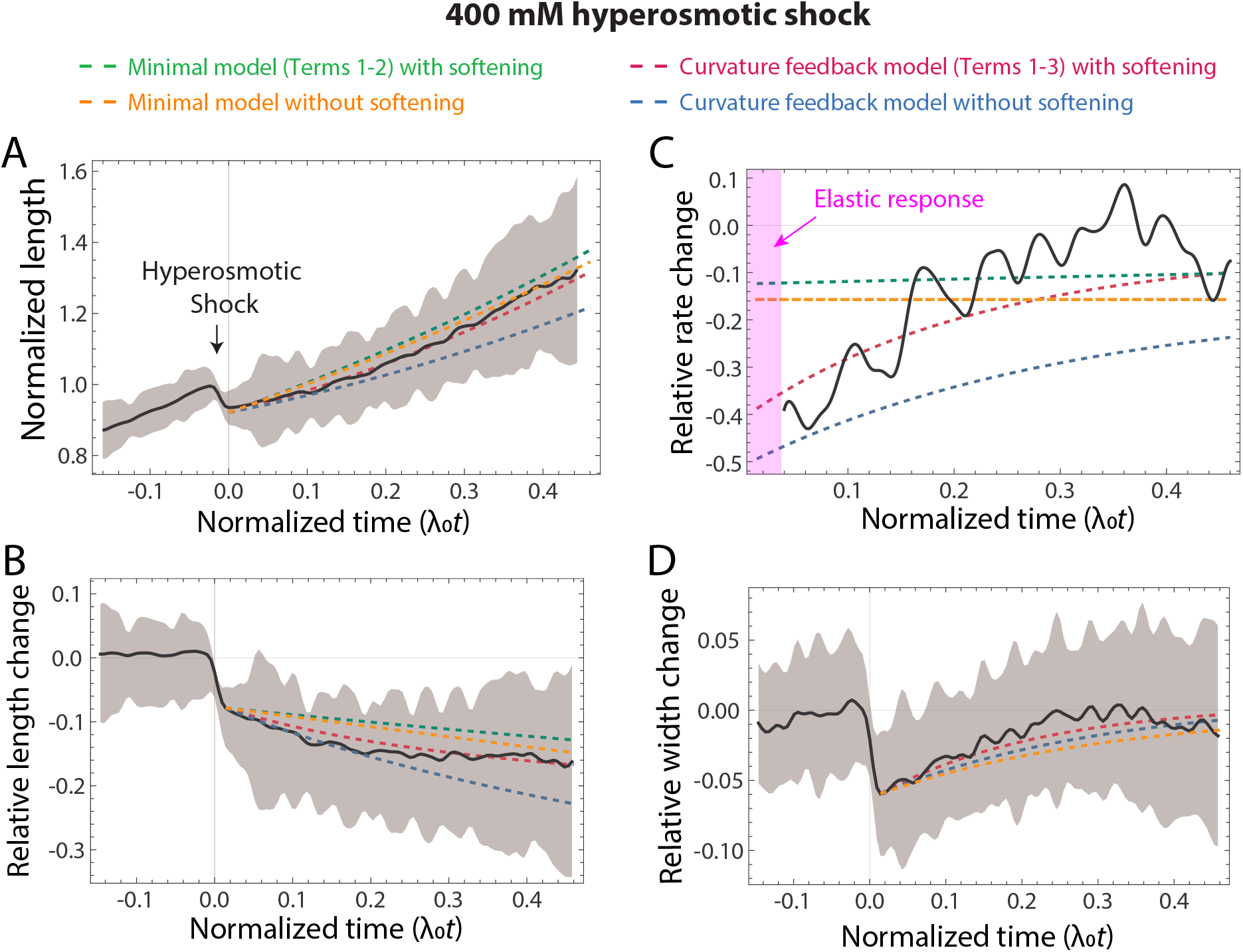
Model with envelope softening predicts homeostasis of growth rate and cell width in response to hyperosmotic shock. (A) The response dynamics of cell length to a 400-mM hyperosmotic shock predicted by our model with envelope softening and curvature feedback (blue) were in close agreement with experimental measurements (black). Shaded region represents 1 standard deviation. Parameter estimates were obtained as described in the Methods. The predictions from the model without curvature feedback and without envelope softening are shown for comparison. (B) The relative cell length change dynamics for the data in (A) are in close agreement with experimental measurements. The initial decrease is *δL*(0) = −0.08. The predictions from the model without curvature feedback and without envelope softening are shown for comparison. (C) The rapid recovery of growth rate to its pre-shock value (black) is predicted by our curvature feedback model with envelope softening. The initial interval just after the shock (pink) was dominated by the elastic response. (D) Width initially shrank elastically after the hyperosmotic shock due to the decrease in turgor pressure, but then recovered back to its pre-shock value within approximately one cell doubling. The transient dynamics and recovery are due to the presence of curvature coupling in our model.

The role of softening in elongation-rate homeostasis is straightforward to understand mechanistically. First, the reduction in turgor pressure caused by the hyperosmotic shock causes the cell to contract. In our curvature feedback model, this decrease in pressure leads to a reduced elongation rate (directly after the shock) due to contributions from term 1 (Eq. 1), as the contraction reduces the available amount of surface area into which new material is being inserted for a given rest state, and term 3, as elongation rate responds to the turgor-induced decrease in width. As elongation rate decreases, however, and the relative length change increases in magnitude, softening of the envelope allows the reduced turgor pressure to adaptively stretch the envelope to the extent that it was stretched before the shock, thereby re-establishing the steady-state elongation rate. In the minimal model, softening also leads to growth-rate adaptation, but on a much longer time scale (Fig. 3C).

We identified further evidence for our curvature feedback model by comparing its predictions to experimental cell width dynamics. Hyperosmotic shock caused a rapid reduction in width (Fig. 3D), followed by a rapid recovery (Fig. 3D) on the same time scale as elongation rate recovery (Fig. 3C). Using the same set of model parameters as was used to fit elongation rate and length dynamics (Fig. 3B,C), both our curvature feedback model and the minimal model predicted this qualitative width recovery (with or without softening) (Fig. 3D). Both softening and curvature feedback were predicted to accelerate width recovery, and the combination of both led to close agreement with experimental measurements (Fig. 3D).

### Softening-mediated elongation-rate homeostasis is unstable for large hyperosmotic shocks

Thus far, we have focused on scenarios in which turgor pressure remains positive after the hyperosmotic shock. Within our model, if a hyperosmotic shock is large enough to cause turgor pressure to fall below zero, softening combined with the compressive forces in the cell envelope due to negative turgor pressure would lead to growth arrest rather than promoting recovery of elongation rate. Mathematically, this feedback manifests as an instability for large-magnitude shocks (Fig. 4A). At long times, our curvature feedback model with softening yields

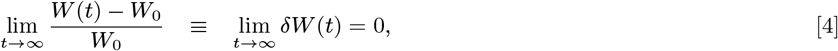

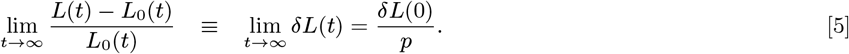

**Fig. 4.**
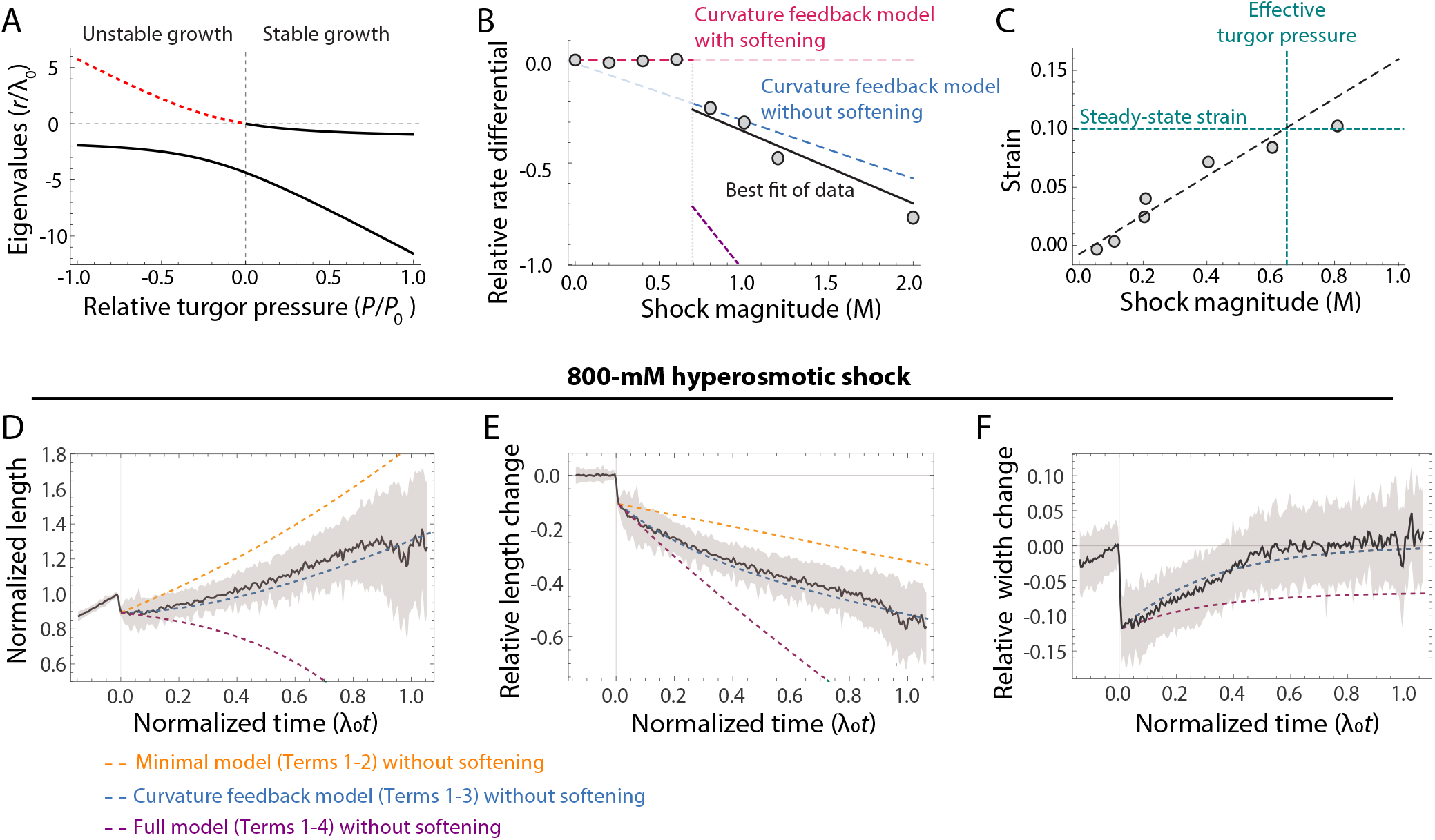
Envelope softening results in an instability for large hyperosmotic shocks. (A) Eigenvalues that dictate the growth or decay of modes of Eq. 10 and 11. When *p* < 0, one eigenvalue becomes positive, indicating an unstable mode. (B) For hyperosmotic shocks below ~ 700 mM in magnitude, cells recovered to their pre-shock growth rate *λ*_0_. For shocks > 700 mM, cells equilibrated to a new growth rate λ *f* that decreased with shock magnitude (Fig. S4). The relative rate change is (*λ_f_* – *λ*_0_)/*λ*_0_. The dashed lines show our model prediction with envelope softening (for shocks < 700 mM) and without (for shocks > 700 mM), which is in close agreement with the experimental data (gray circles). The purple line represents the prediction from the model in the presence of strain sensing (*g*_3_ = 2). (C) Magnitude of the elastic length strain after a hyperosmotic shock of various magnitudes. The length strain reaches the value *ε*_0_ =0.1 (the estimated envelope strain during steady-state growth) at an estimated shock magnitude of ~ 700 mM. (D) After the 800-mM hyperosmotic shock, length increased much more slowly than the predicted rate for an unperturbed cell. Model predictions with fixed Young’s modulus (blue) are in close agreement with experimental data (black). (E) Since growth rate decreased after the shock and remained below its unperturbed value, the relative length change *δL*(*t*) continued to decrease from its initial value of −0.11 directly after the shock to an asymptote of −1. In the absence of curvature feedback (orange), the same final growth rate is attained as with curvature sensing, but the predicted transient behavior does not agree with experimental data (black), providing evidence that length is coupled to width through curvature feedback. (F) Cell width recovers to its pre-shock value after an 800-mM shock. The close fit of our model (blue dashed curve) to experimental data (black) provides an estimates of the model parameters *g*_1_ =3.2 and *g*_3_ = 0. In the presence of strain sensing (*g*_3_ = 2), our model predicts that width would not recover to its pre-shock value (purple).

That is, width recovers to its pre-shock steady-state value, while the relative difference between cell length and the length prescribed by steady-state elongation diverges as *p* → 0.

Based on this analysis, we hypothesized that cells abandon softening after large hyperosmotic shocks and thus we explored this regime experimentally. Remarkably, the relative difference between the steady-state elongation before and after hyperosmotic shock exhibited a sharp discontinuity at a critical shock magnitude (Fig. 4B, S4). For all shocks < 700 mM, elongation rate recovered to its unperturbed value (consistent with envelope softening). Conversely, for larger shocks, cells continued to grow (in contradiction to model predictions with envelope softening), but eventually stabilized at an elongation rate that decreased linearly with shock magnitude (Fig. 4B). This behavior was in quantitative agreement with our theory. Specifically, our Curvature Feedback model without softening predicts that the slope of the linear dependence between elongation rate and shock magnitude is twice the slope of envelope strain as a function of shock magnitude (Methods). This model prediction of ≈ 2 * (−0.14) = −0.28 (Fig. 4B-C) is in reasonable agreement with the experimental slope of −0.35 ± 0.07. According to our theory, this linear decrease arises because of indirect mechanical effects of turgor pressure on elongation rate via cell envelope geometry. That is, reduction in turgor pressure causes a decrease in cell surface area, which reduces elongation rate by reducing the amount of cell envelope synthesis via Term 1 in Eq. 1.

For shocks with magnitude above the 700 mM threshold, direct feedback between envelope curvature and envelope expansion (Eq. 1, Term 3) was critical for explaining the dynamics of cell length. The minimal model with only Terms 1 and 2 failed to predict the transient length and elongation dynamics after the shock, which were precisely explained by our curvature feedback model (Fig. 4D-E, S5). Together, these data suggest that envelope softening and stored growth are abandoned for large magnitude shocks, and provide further support that envelope expansion is directly dependent on envelope curvature.

What is the origin of the 700 mM threshold? Within our model, this threshold is simply the steady-state turgor pressure (in units of osmolarity). Previous studies using atomic force microscopy (AFM) estimated the turgor pressure in *E. coli* to be ≈ 50 – 150 mM (30), consistent with our observation that cells plasmolyze for shock magnitudes ≥ 100 mM (3). However, we previously identified a surprising and interesting caveat to these measurements: the cell envelope clearly continues to contract linearly for shock magnitudes up to 700 mM (Fig. 4C) (3). In other words, even though turgor pressure is zero after shocks > 100 mM, the cytoplasm still induces tensile stress in the envelope unless the shock is > 700 mM; the degree of contraction was maintained when the shock magnitude was increased from 700 mM to 3 M (Fig. 4C). Since this tensile stress, and not turgor pressure itself, is the key variable that is relevant to cell-envelope expansion, we define the effective turgor pressure as 700 mM, which precisely agrees with the critical shock magnitude above which we observe the discontinuity in steady-state growth rate (Fig. 4B) and predict that cell-envelope softening is abolished.

### Envelope expansion is not explicitly dependent on mechanical strain

In all the above analyses, it was unnecessary to invoke an explicit dependence of envelope expansion on envelope strain (Eq. 1, Term 4). In fact, with inclusion of this dependence our model predictions are inconsistent with experimental measurements (Fig. 4B-F). Under explicit strain-dependent envelope expansion, altering pressure shifts the steady-state width, which thus recovers to a different value after the shock compared with the pre-shock width (Fig. 4F, Methods). Furthermore, inclusion of strain-dependent expansion led to poor predictions of the elongation rate dynamics for shock magnitudes above the critical threshold (Fig. 4B, Methods). In contrast, the successful prediction of both these behaviors by our model in the absence of Term 4 in Eq. 1 provides support for the curvature feedback model without explicit strain dependence.

## Discussion

By combining theory and experiment, we identified two functional relationships between key cell-scale variables that promote homeostasis during cell morphogenesis: i) softening of the cell envelope due to mismatch between elongation rate and envelope-synthesis rate, and ii) the direct dependence of cell elongation rate on envelope curvature (“curvature feedback”). These two relationships are both required to guarantee the experimentally observed homeostasis in cell growth rate (Fig. 3A-C), while only curvature feedback is required to guarantee cell-width homeostasis (Fig. 3D, 4F), although softening does accelerate width recovery after a shock (Fig. 3D). Our complete model (Eq. 1-3) suggests that *E. coli* morphogenesis relies on interlocking negative feedback loops linking three key cellular-scale variables: cell width, cell length, and envelope stiffness (Fig. 5A). Negative feedback has been demonstrated to increase stability and accelerate the response of control systems (31), underscoring the likely fitness advantages of homeostatic mechanisms during cell morphogenesis.

**Fig. 5.**
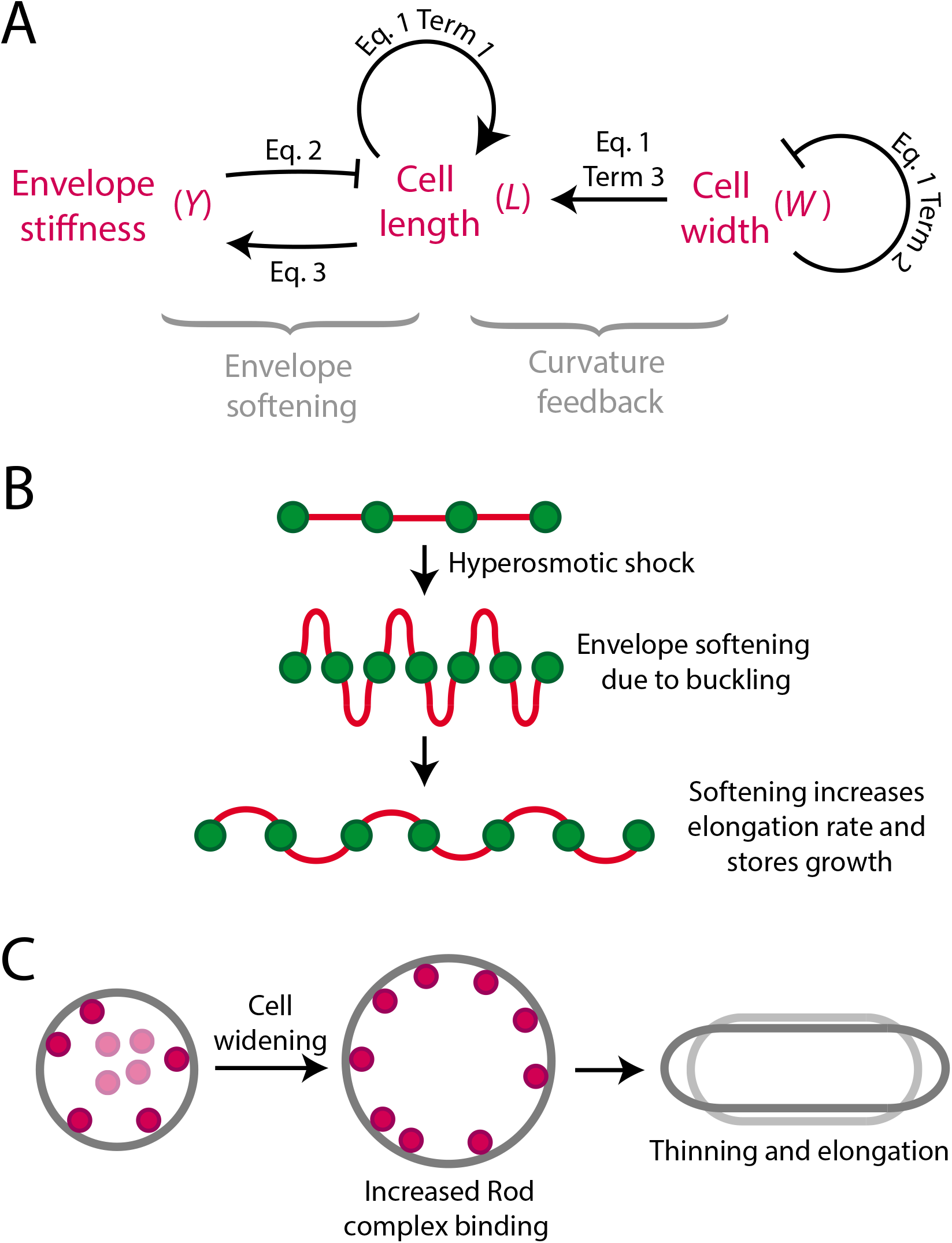
Feedback linking cell length to envelope stiffness cell width leads to growth rate and width homeostasis. (A) Summary of model findings. During steady state growth, cell length increases exponentially due to Term 1 in Eq. 1, and Term 2 ensures width homeostasis. After a hyperosmotic shock is applied, pressure is reduced, which affects length and width through elastic stretching as well as through coupling between width and length through Term 3 in Eq. 1. Envelope stiffness inhibits elongation through Eq. 2, and a hyperosmotic shock induces softening through Eq. 3. The softening mechanism is assumed to only be active when the effective turgor pressure is positive to avoid the predicted instability (Figure 4A). Softening compensates for the decrease in width and growth rate after the shock and leads to growth rate homeostasis. (B) Toy model of softening mechanism. After a hyperosmotic shock, newly inserted cell-wall material could be wrinkled, hence would be softer than fully stretched material and thereby reduce the cell envelope Young’s modulus. (C) Toy model illustrating curvature feedback. Enzymes responsible for envelope synthesis may prefer binding to certain local curvatures, such that cell widening promotes increased activity that leads to simultaneous thinning and elongation.

In our model, the interactions between cell-scale morphogenetic variables (Eq. 1, Fig. 5A) must be mediated by molecular-scale properties such as cell envelope synthesis and envelope curvature. Our results point to potential molecular mechanisms that underlie these interactions. First, we previously speculated that molecular scale wrinkling of the cell envelope could account for stored growth (3). In this picture, which is consistent with our theory of envelope softening, hyperosmotic shock reduces elongation rate more than it reduces the rate of cell envelope synthesis. As a result, new cell envelope synthesis wrinkles as it is inserted into the cell wall (Fig. 5B). This wrinkled material results in envelope softening since tensile forces will tend to unwrinkle it prior to stretching. Second, one plausible mechanism for feedback between envelope curvature and envelope expansion (Eq. 1, Terms 2 and 3) would be curvature-dependent localization of the Rod complexes. Rod complexes play a key role in cell elongation by coordinating peptidoglycan synthesis, and alteration of their activity in Bacillus subtilis changes cell width (20) via an unknown mechanism. Our model suggests that if the binding affinity of Rod complexes increases as envelope curvature decreases, wider cells would tend to simultaneously thin and elongate faster (Fig. 5C).

An interesting finding in our study is the disparity between the hyperosmotic shock magnitude at which cells plasmolyze (~ 100 mM) and that at which the cell envelope reaches its rest state (~ 700 mM). We speculate that this disparity is a direct result of two cellular features: the stiffness of the cytoplasm and the strong mechanical connections between the cytoplasmic (inner) membrane and the cell envelope. If the cytoplasm is both relatively stiff and connected to the cell envelope, then hyperosmotic shocks between 100 mM and 700 mM could cause compression of the cytoplasm/cell envelope complex. In this case, the inner membrane would be balancing tensile forces in the cell envelope with shear forces between the two structures. This question should be an interesting topic for future study.

It remains unclear what prevents cells from implementing envelope softening for high-magnitude shocks. For such shocks, the substantial plasmolysis (separation of the cytoplasm from the cell wall) may inhibit wall synthesis by preventing the wall synthesis enzymes from reaching from the inner membrane to the cell wall. Future experiments that can localize the synthesis machinery with super-resolution during osmotic perturbations may resolve this mystery.

Homeostasis of cellular morphogenesis is conserved in most (if not all) cells. The systems that guarantee this homeostasis must inherently incorporate molecular-scale and cell-scale information. Here, we provided a paradigm for this integration for one of the best-studied model organisms. One high-level takeaway from our analysis is that this homeostatic system is complex: maintenance of cell width and elongation rate cannot be decomposed into component systems but are intimately coupled (Fig. 5A). It will be interesting to compare parallel systems in other organisms with other cellular morphologies to test the generality of this architecture.

## Materials and Methods

### Model for growth without envelope softening

Approximating cell shape using a cylinder whose width *W*(*t*) and length *L*(*t*) change with time, we can express the position of a point, parameterized by *z* and *θ* (Fig. 1), as

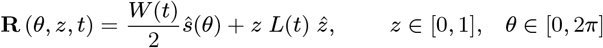

where *ŝ*(*θ*) and 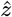 are cylindrical basis vectors. This cylindrical ansatz, with *W*(*t*) = *W*_0_ and *L*(*t*) = *L*_0_(*t*), is a solution of Eq. 1 (SI), which we interpret as the steady-state behavior of *E. coli*. When cells are perturbed away from the steady-state behavior of Eq. 1, in the absence of softening and to linear order in deviation, the width and length behave as

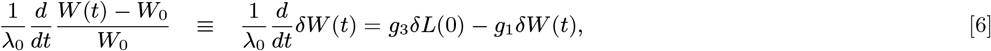

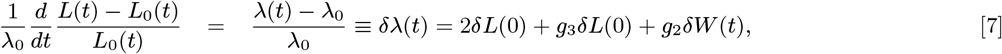

where *g*_1_ = (*α*_1_ – *α*_2_ – 4)/4, *g*_2_ = *α*_2_/4, and *g*_3_ = *β*/2 (SI).

The absence of *δL*(*t*) from the right hand side of Eq. 6-7 can be understood since the change in length does not manifest as a change in curvature or strain, which are local quantities. As a result, growth rate does not recover in our model in the absence of softening.

At long times (steady state) the solution to Eq. 6-7 is given as

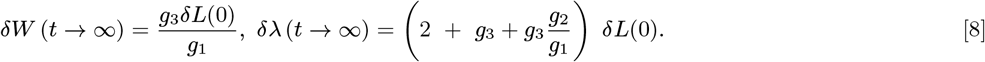

This results shows that width does not recover to the steady state value in the presence of the direct strain sensing term (*g*_3_ ≠ 0) and the relative change in elongation rate will not approach 2*δL*(0).

### Model for growth with envelope softening under the stored growth constraint

During the period of low turgor pressure after the hyperosmotic shock, the length of the cell *L*(*t*) dictated by Eq. 2 is lower than that of an unperturbed cell, *L*_0_(*t*). The stored growth condition demands that these two lengths are equal when pressure is restored (*p* = 1), which leads to the relation

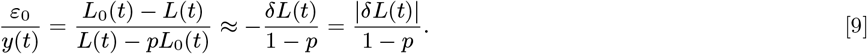

Thus, the value of normalized Young’s modulus *y*(*t*) = *Y*(*t*)/*Y*_0_ becomes a proxy for the length deviation *δL*(*t*).

Using the relation above (Eq. 9) in Eq. 1, to linear order in length deviation and width deviation *δW*(*t*) we obtain

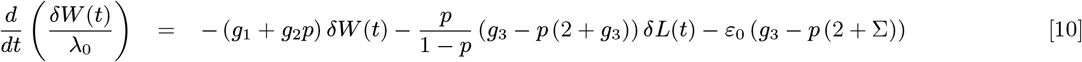

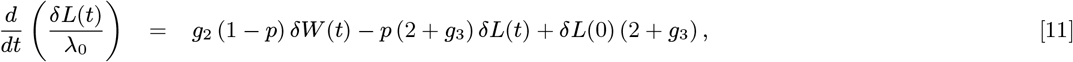

Interestingly, cell envelope softening both leads to growth rate homeostasis (*δL*(*t* → ∞) = constant) and enhances width homeostasis: width recovers at the faster rate (*g*_1_ + *g*_2_*p*), compared with *g*_1_ in the case of fixed Young’s modulus (Eq. 6). This fact is consistent with our observation that the rate of width recovery after a 400-mM shock (Fig. 2D) is faster than after an 800-mM shock (Fig. 4F).

By setting the left-hand sides of Eq. 10 and 11 to zero, we find that the solutions for length and width deviation should approach

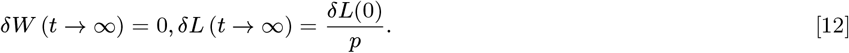

Note that as p approaches 0, the deviation in length increases and the equations lead to an instability when *p* ≤ 0.

### Data analysis

Each curve in Fig. 2 – 4 and S2 - S5 was obtained by averaging over all single-cell traces after normalizing by the value at the time point immediately before the hyperosmotic shock. The unperturbed growth rate and cellular dimensions were determined from the period before the shock, from which we extrapolated the growth function *L*_0_(*t*). *δL*(0) and *δW*(0) were computed from the decrease in length and width in the 25 s directly after the shock, from which we obtained the data shown in Fig. 4C.

The parameters that are needed to determine the behavior of width and length in our model are *g*_1_,*g*_2_,*g*_3_, and *ε*_0_. To fit our model to experimental data, we take *ε*_0_ ≈ 0.1 from previous measurements (5) and minimize the error function

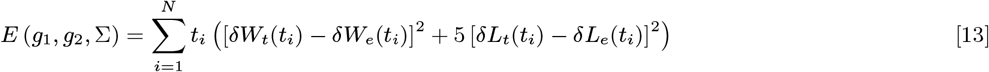

over data points *i* and the subscripts *t* and *e* refer to theoretical prediction and experimental measurement, respectively. We weighted later times more heavily since the initial conditions of the model already match the experimental data. We also weighted the length measurements 5-fold more heavily since it depends on all parameters and is coupled to width, in contrast width does not depend on the value of *g*_2_ or length (Eq. 6-7). By minimizing this error function using simulated annealing with data for an 800-mM hyperosmotic shock, we obtained estimates of the parameters *g*_1_ = 4.4, *g*_2_ = 7.0, and *g*_3_ = 0.23.

To assess the robustness of these values, we examined the eigenvalues and eigenvectors of the Hessian of *E*(*g*_1_, *g*_2_, *g*_3_) near this minimum. Directions in parameter space with higher eigenvalues are expected to be more tightly constrained by the data. The eigenvalues of the Hessian (normalized by the minimum value of the function) were 164, 0.4, and 0.05, with corresponding eigenvectors *v*_1_ = (−0.21, 0.16, 0.96), *v*_2_ = (0.44, −0.86, 0.24), and *v*_3_ = (0.87, 0.48, 0.11).

We rewrote the parameters in terms of coordinates *G*_1_,*G*_2_,*G*_3_ along the eigenvectors *v*_1_,*v*_2_, and *v*_3_ as *g*_1_ = 4.4 – 0.21*G*_1_ + 0.44*G*_2_ + 0.87*G*_3_, *g*_2_ = 7.0 + 0.16*G*_1_ – 0.86*G*_2_ + 0.48*G*_3_, and *g*_3_ = 0.23 + 0.96*G*_1_ + 0.24*G*_2_ + 0.11*G*_3_. By varying the parameters *G_i_* individually, we obtained confidence intervals for these parameters.

In addition to fitting the parameters by the optimization procedure described above, we systematically varied the parameter values and verified the quality of the fit. Nearly all experimental results in this study were well fit by the values *g*_1_ = 4.4, *g*_2_ = 7.0, and *g*_3_ = 0.23. For the width dynamics data, *g*_2_ appeared slightly overestimated, with *g*_2_ = 6.0 providing a slightly better fit. We also obtained fits of *g*_1_ = 3.2, *g*_2_ = 5.7, from the curvature-coupled model in the absence of strain coupling (*g*_2_ = 0), approximately consistent with our other fits. In the minimal model (*g*_2_ = 0, *g*_3_ = 0) the value of *g*_1_ = 3.2, is the same as it would be in the curvature feedback model since *g*_2_ does not influence width dynamics.

## ACKNOWLEDGMENTS

The authors thank the Huang and Rojas labs for useful discussions. K.C.H. is a Chan Zuckerberg Biohub Investigator. K.C.H. was supported in part by the NSF grant EF-2125383. A.G was supported in part by the National Science Foundation under Grant DMS-1616926, the NSF-CREST Center for Cellular and Bio-molecular Machines at UC Merced (HRD-1547848) and the NSF Center for Engineering Mechanobiology grant (CMMI-154857). This work was also supported in part by the National Science Foundation under Grant PHYS-1066293 and the hospitality of the Aspen Center for Physics.

