## Supplementary Information for "Feedback linking cell envelope stiffness, curvature, and synthesis enables robust rod-shaped bacterial growth"

#### This PDF file includes:

Supplementary text
Figs. S1 to S5
SI References

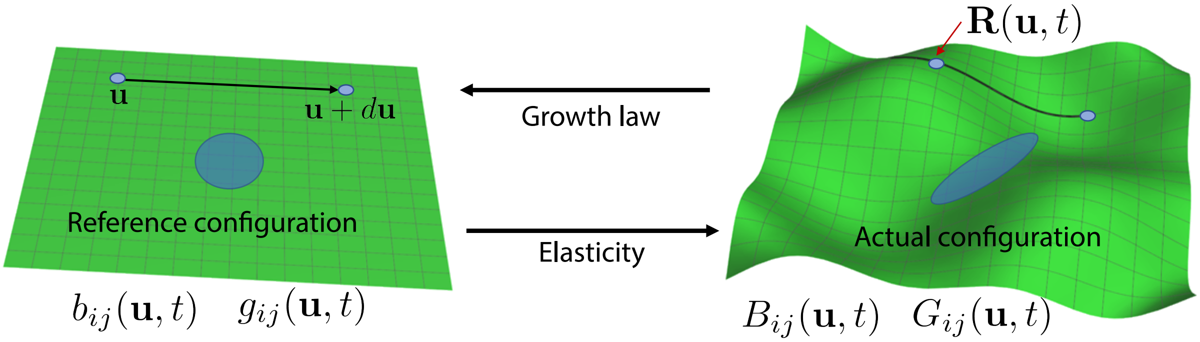


**Fig. S1. Schematic of growth kinematics and dynamics.** A surface is labeled by coordinates $\mathbf{u} \equiv (u_1, u_2)$ and the positions of these points is given by the function $\mathbf{R}(\mathbf{u}, t)$. The metric tensor, $G_{ij}(\mathbf{u}, t)$, is used to calculate infinitesimal lengths of segments between $\mathbf{u}$ and $\mathbf{u} + d\mathbf{u}$. The curvature tensor, $B_{ij}(\mathbf{u}, t)$, described normal curvature of the curve tangent to the direction $d\mathbf{u}$ at $\mathbf{u}$. The rest metric tensor $g_{ij}(\mathbf{u}, t)$ represents the unstretched lengths of these curves and the rest curvature tensor $b_{ij}(\mathbf{u}, t)$ represents their unbent curvature. The blue ellipse on the right is the actual configuration of the circle shown in the reference configuration on the left. Black arrows represent the direction of influence during growth: geometry affects growth through feedback, which in turn changes the geometry through elasticity.

### Supporting Information Text

**Kinematics and dynamics of growing surfaces.** We describe a surface embedded in 3D space by parameterizing the points on it with a coordinate system $\mathbf{u} \equiv (u_1, u_2)$ and giving the position of each point, at time $t$, as $\mathbf{R}(\mathbf{u}, t)$. We assume that the coordinates $(u_1, u_2)$ are fixed on the surface over time so that $\mathbf{R}(\mathbf{u}, t)$ gives the trajectory of a labeled point on the surface (cell envelope) over time. To describe the length between two nearby points $\mathbf{u}$ and $\mathbf{u} + d\mathbf{u}$, we use the chain rule, which gives,

$$d\mathbf{R}(u_1, u_2, t) \cdot d\mathbf{R}(u_1, u_2, t) = \sum_{i,j=1}^{2} \partial_i \mathbf{R} \cdot \partial_j \mathbf{R}\, du_i\, du_j \equiv \sum_{i,j=1}^{2} G_{ij}(u_1, u_2, t)\, du_i\, du_j, \qquad \text{[S.1]}$$

where $\partial_j$ represents partial differentiation with respect to $u_i$ and we have defined the metric tensor $G_{ij}$. Therefore, the metric defines lengths on the surface and the rest metric $g_{ij}$ defines the corresponding rest lengths that are preferred by the elastic energy. Raising the indices of the metrics, $G^{ij}$ and $g^{ij}$, denotes the corresponding inverse. The curvature tensor is defined as $B_{ij}(\mathbf{u}, t) = \hat{\mathbf{n}} \cdot \partial_i \partial_j \mathbf{R}$, where $\hat{\mathbf{n}}(\mathbf{u}, t)$ is the unit normal to the surface at each point, and gives the curvature of curves in the direction normal to the surface. The rest curvature tensor $b_{ij}(\mathbf{u}, t)$ gives the values preferred by the elastic energy. We will assume in the following that the rest curvature tensor vanishes, $b_{ij}(\mathbf{u}, t) = 0$. While this simplifying assumption will not change our conclusions, biologically it means that if we cut open the cell envelope it will flatten its shape.

We take advantage of the large separation between growth ($\sim 10$ min) and envelope elastic ($\ll 1$ s) time scales to separate conceptually the growth process from the elastic response of the cell to changes in applied forces. The elastic response of the cell envelope can be modeled as a thin elastic shell whose energy is quadratic in the strain (1):

$$\mathcal{E} = \frac{h}{8} \int d^2\mathbf{u} \sqrt{g} \sum_{i,j,k,\ell=1}^{2} A^{ijk\ell} \left[ (G_{ij} - g_{ij})\,(G_{k\ell} - g_{k\ell}) + \frac{h^2}{2}\,(B_{ij} - b_{ij})\,(B_{k\ell} - b_{k\ell}) \right], \qquad \text{[S.2]}$$

where $h$ is the thickness of the cell envelope and $A^{ijk\ell}$ is the elasticity tensor, which for isotropic materials is given by $A^{ijk\ell} = \lambda\, g^{ij} g^{k\ell} + 2\,\mu\, g^{ik} g^{j\ell}$. Here $\lambda$ and $\mu$ are the (2D) Lamé constants, related to Young's modulus $E$ and Poisson's ratio $\nu$ through $\lambda = E\nu/(1-\nu^2)$ and $2\mu = E/(1+\nu)$. Note that the elastic strain can be defined as the difference between the metrics, $S_{ij} \equiv (G_{ij} - g_{ij})/2$.

The quantities $A^{ijk\ell}(\mathbf{u}, t)$, $g^{k\ell}(\mathbf{u}, t)$, and $b^{k\ell}(\mathbf{u}, t)$ define the elastic behavior of the cell envelope, which in turn gives the shape of the envelope $\mathbf{R}(\mathbf{u}, t)$ through minimization of the energy. Thus, to determine how the system evolves over time in response to perturbations we need to know how $A^{ijk\ell}(\mathbf{u}, t)$ and $g^{k\ell}(\mathbf{u}, t)$ are biologically regulated.

**Growth and feedback laws.** We construct a minimal model capable of capturing the mechanical behavior of the cell envelope (described in the previous section) and the regulation of its properties during growth. Intrinsic changes in cell shape, which are due to insertion of new material rather than changes in applied forces, can quantified as changes in rest lengths between points in the cell envelope and, therefore, are mathematically described by the time derivative of the rest metric $g^{k\ell}(\mathbf{u}, t)$. Experimentally, it was observed that growth rate depends on both envelope curvature (through the action of MreB) and strain (2–4).

To derive a minimal growth law, and in agreement with experiments (3), we assume that this dependence is local: Growth, at a given point in the cell envelope, depends smoothly on the shape and strain in the immediate vicinity of that point. We derived in Ref. (5) a generic growth law consistent with these assumption as a Taylor expansion in curvature and strain. To second order in curvature and linear order in strain we have:

$$\frac{1}{\lambda_0}\partial_t g_{ij} = \sigma_1 G_{ij} - \frac{\sigma_2}{H_0}B_{ij} + \frac{\alpha_1}{2H_0^2}(H-H_0)B_{ij} + \frac{\alpha_2}{2H_0}(H_0-H)G_{ij} + \beta_1 S_{ij} + \beta_2 S\, G_{ij}$$
$$\frac{\gamma}{(H_0)^2}K\, G_{ij} + \frac{\delta}{4(H_0)^2}(H_0-H)^2 G_{ij}, \qquad \text{[S.3]}$$

where $H(\mathbf{u},t) \equiv G^{ij}(\mathbf{u},t)B_{ij}(\mathbf{u},t)$ is twice the mean curvature, $K(\mathbf{u},t)$ is the Gaussian curvature, and $S(\mathbf{u},t) \equiv G^{ij}(\mathbf{u},t)S_{ij}(\mathbf{u},t)$ is the areal strain. Here $\lambda_0$ gives an inverse time scale that we chose as the steady-state growth rate, $W_0 = 2/H_0$ gives a reference length scale that we chose as the steady-state width of the cell. The parameters $\alpha_i, \beta_i, \sigma_i, \gamma$, and $\delta$ are dimensionless numbers characterizing the feedback law. The first two terms, proportional to $\sigma_1$ and $\sigma_2$ are the terms we grouped into $A_{ij}(\mathbf{u},t)$ in the main text and $\beta = \beta_1$.

Note that, without loss of generality, we wrote Eq. (S.3) so that all but the first two shape-dependent terms vanish at steady state. Parameterizing the growth rate in this way justifies excluding higher order terms from our calculations, since these will be subdominant for small deviations from the steady state width and small strains (which is the regime explored in our experiments). Additionally, we do not expect higher order derivatives of curvature, such as the Laplacian of mean curvature ($\triangle H$) for example, to appear in the growth law since curvature sensing proteins, such as MreB, are small compared with the cell envelope size and sensing these higher order terms will be more difficult.

**Cylindrical solutions.** We next focus on cylindrical solutions to the above equations. A cylinder with time dependent width $W(t)$ and length $L(t)$ can be written as

$$\mathbf{R}(\theta,z,t) = \frac{W(t)}{2}\hat{s}(\theta) + z\,L(t)\,\hat{z}, \qquad z\in[0,1], \quad \theta\in[0,2\pi), \qquad \text{[S.4]}$$

where $\hat{s}$ and $\hat{z}$ are cylindrical basis vectors. Note again that this coordinate system is adapted to the growing cylinder so that $\mathbf{R}(\theta,z,t)$ gives the trajectory of a labeled point, with fixed coordinates $(\theta,z)$, over time. The metric, rest metric, and curvature tensors are given by

$$G_{ij}(\theta,z,t) = \begin{pmatrix} W^2(t)/4 & 0 \\ 0 & L^2(t)\end{pmatrix}, \qquad g_{ij}(\theta,z,t) = \begin{pmatrix} w^2(t)/4 & 0 \\ 0 & l^2(t)\end{pmatrix}, \qquad B_{ij}(\theta,z,t) = \begin{pmatrix} W(t)/2 & 0 \\ 0 & 0\end{pmatrix} \qquad \text{[S.5]}$$

If we know the rest length and width, in addition to the elastic moduli of the cell envelope, we can solve for the actual length and width by using the equations of elastic equilibrium. Plugging in Eq. (S.4) into the elastic energy Eq. (S.2) and minimizing with respect to $L(t)$ and $W(t)$ gives the thin pressure vessel equations:

$$W(t) = w(t)\left[1 + \frac{P(t)w(t)}{2hY_\theta(t)}\left(1+\frac{\nu}{2}\right)\right], \qquad \text{[S.6]}$$
$$L(t) = l(t)\left[1 + \frac{P(t)w(t)}{2hY_z(t)}\left(\frac{1}{2}+\nu\right)\right], \qquad \text{[S.7]}$$

where $P$ is the turgor pressure, $h$ is the thickness of the shell, and $\nu$ is the Poisson ratio. We allow for the possibility of different Young's moduli in the $\theta$ and z directions (Yao et al., 1999). Using estimates of these parameters from (Deng et al., 2011; Tuson et al., 2012), $Y_z = 23 \pm 8$ MPa and $Y_\theta = 49 \pm 20$ MPa, $P_0 = 29 \pm 3$ kPa, $h = 4.5 \pm 1.5$nm, and $W_0 = 1.1\mu$m. For $\nu = 0$, $\varepsilon_\theta \equiv (W(t)-w(t))/w(t) \approx \varepsilon_z = (L(t)-l(t))/l(t) \approx 0.07$. These values are consistent with our experimental observations that the elastic strain in $L(t)$ and $W(t)$ due to a hyperosmotic shock are similar. Therefore, we assumed $\varepsilon_\theta \approx \varepsilon_z \approx \varepsilon_0 \approx 0.1$ in our calculations, where the last approximate equality comes from experiments (Rojas et al. 2014) as described in the results section. We also assume that the relative change in Young's modulus is the same in both direction so that $y(t) = Y_z(t)/Y_z(0) = Y_\theta(t)/Y_\theta(0)$, where the value at $t = 0$ is the normal steady state value. Equations (S.6-S.7) can now be written as

$$W(t) = w(t)\left[1 + \epsilon_0\frac{p(t)}{y(t)}\right] \equiv w(t)\left[1+\epsilon_0 u(t)\right], \qquad \text{[S.8]}$$
$$L(t) = l(t)\left[1 + \epsilon_0\frac{p(t)}{y(t)}\right] \equiv l(t)\left[1+\epsilon_0 u(t)\right], \qquad \text{[S.9]}$$

where $p(t) = P(t)/P_0$. Note that pressure and Young's modulus enter together into all expression in the combination $u(t) = p(t)/y(t)$.

We can obtain the equations of motion for the rest length and width by plugging Eqs. (S.4-S.5) into the growth law Eq. (S.3) to get

$$\frac{1}{\lambda_0}\frac{\partial_t w^2(t)}{W^2(t)} = \sigma_1 - \sigma_2\frac{W_0}{W} - \frac{\alpha_1}{2}\left(1-\frac{W_0}{W}\right)\frac{W_0}{W} + \frac{\alpha_2}{2}\left(1-\frac{W_0}{W}\right) + (\beta_1+\beta_2)\frac{W^2-w^2}{2W^2} + \beta_2\frac{L^2-l^2}{2L^2} + \frac{\delta}{4}\left(1-\frac{W_0}{W}\right)^2, \qquad \text{[S.10]}$$
$$\frac{1}{\lambda_0}\frac{\partial_t l^2(t)}{L^2(t)} = \sigma_1 + \frac{\alpha_2}{2}\left(1-\frac{W_0}{W}\right) + (\beta_1+\beta_2)\frac{L^2-l^2}{2L^2} + \beta_2\frac{W^2-w^2}{2W^2} + \frac{\delta}{4}\left(1-\frac{W_0}{W}\right)^2. \qquad \text{[S.11]}$$

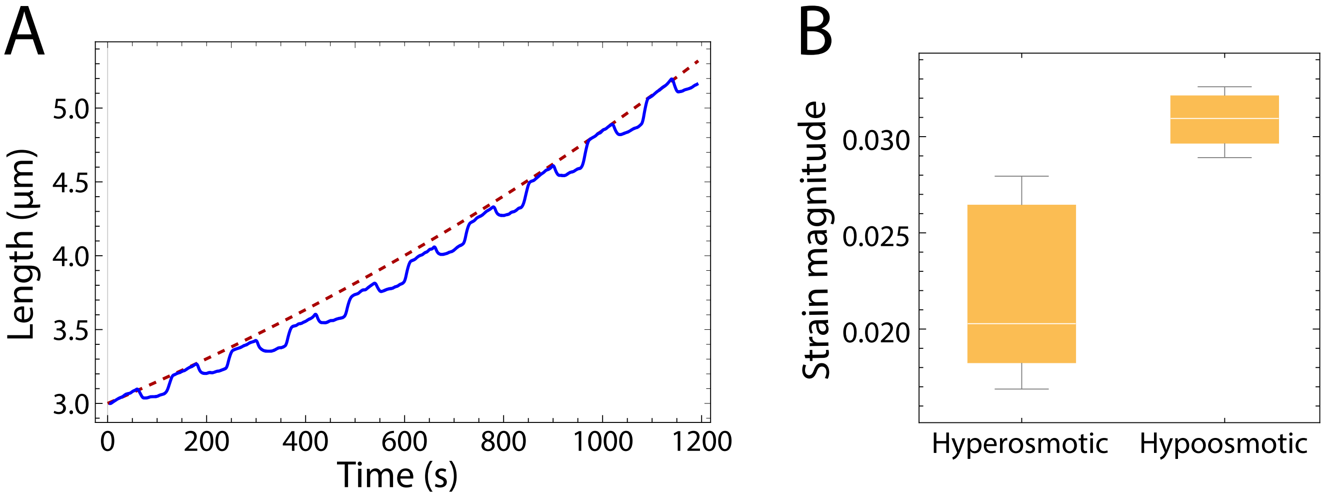


**Fig. S2. Cyclic osmotic shocks of magnitude 100-mM with period 120 seconds.** (A) Experimentally measured cell length dynamics (blue) in response to cycles of 100-mM hyper- and hypoosmotic shocks. (B) The mechanical strain induced by the hyperosmotic shocks in (A) is smaller than the strain induced by the hypoosmotic shocks, implying softening of the cell envelope.

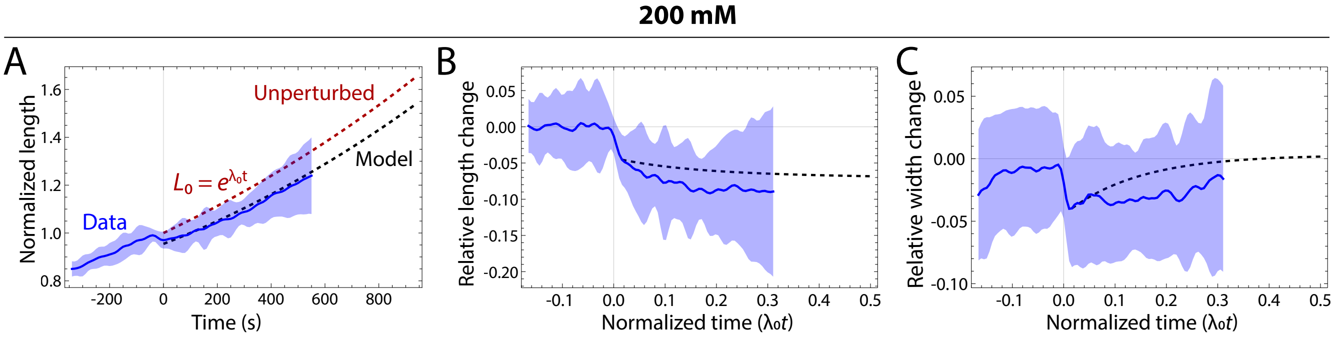


**Fig. S3. Model predicts relative length and width dynamics after a 200-mM hyperosmotic shock.** (A) The predicted dynamics of cell length in response to a 200-mM hyperosmotic shock from our model with envelope softening (black dashed line) were in reasonable agreement with experimental measurements (blue). The red dashed curve represents the extrapolated length of an unperturbed cell based on the pre-shock length and growth rate. Shaded region represents 1 standard deviation. The parameter estimates $g_1 = 3.2, g_2 = 5.7$, and $\varepsilon_0 = 0.1$ were used. (B) The predicted relative length change dynamics calculated from (A) (black dashed line) are in reasonable agreement with experimental measurements (blue). The measured initial decrease directly after the shock was $\delta L(0) = -0.045$. Shaded region represents 1 standard deviation. (C) Width initially shrank elastically after the hyperosmotic shock due to the decrease in turgor pressure, but then recovered back to its pre-shock value within approximately one cell doubling. The transient dynamics and recovery are strongly dependent on the presence of curvature coupling in the model. The measured initial decrease was $\delta W(0) = -0.04$. Shaded region represents 1 standard deviation.

So far, $W_0$ has only played the role of an arbitrary parameter that carries units of length in the growth law. If we chose a different value for it, all that would happen is a redefinition of the growth parameters (such as $\sigma_2$). By choosing $W_0$ to be the steady state width, we obtain that $\sigma_2 = \sigma_1 + (\beta_1 + 2\beta_2)\varepsilon_0$ to linear order in strain. This is not a fine tuning of the parameters of the growth law, but is simply a matter of defining the parameter $W_0$. Similarly, by choosing $\lambda_0$ to be the steady state growth rate, we obtain

$$\sigma_1 = 2 - (\beta_1 + 2\beta_2 + 2)\varepsilon_0 + O(\varepsilon_0^2). \qquad \text{[S.12]}$$

With this choice of parameters, a cylinder with an exponentially expanding length $L(t) = L_0(t) \equiv L(0)\exp(\lambda_0 t)$ and constant width $W(t) = W_0$ is a solution.

In order to predict how the cell will respond as we perturb the steady-state solution, it is useful to define relative width and length changes as $\delta W(t) = (W(t) - W_0)/W_0$ and $\delta L(t) = (L(t) - L_0(t))/L_0(t)$, respectively. The relative width (and to some degree length) change in our experiments is small (below 10%) and we are justified in exploring the equations of motion up to linear order in these quantities. Combining Eqs. (S.8 - S.11) to linear order in $\delta W(t)$ and $\delta L(t)$ gives,

$$\frac{d}{dt}\frac{\delta W(t)}{\lambda_0} = -g_1\,\delta W(t) + (u(t)-1)g_3\varepsilon_0 + \frac{\varepsilon_0}{\lambda_0}u'(t) \qquad \text{[S.13]}$$

$$\frac{d}{dt}\frac{\delta L(t)}{\lambda_0} = g_2\,\delta W(t) + (u(t)-1)(g_3+2)\varepsilon_0 + \frac{\varepsilon_0}{\lambda_0}u'(t), \qquad \text{[S.14]}$$

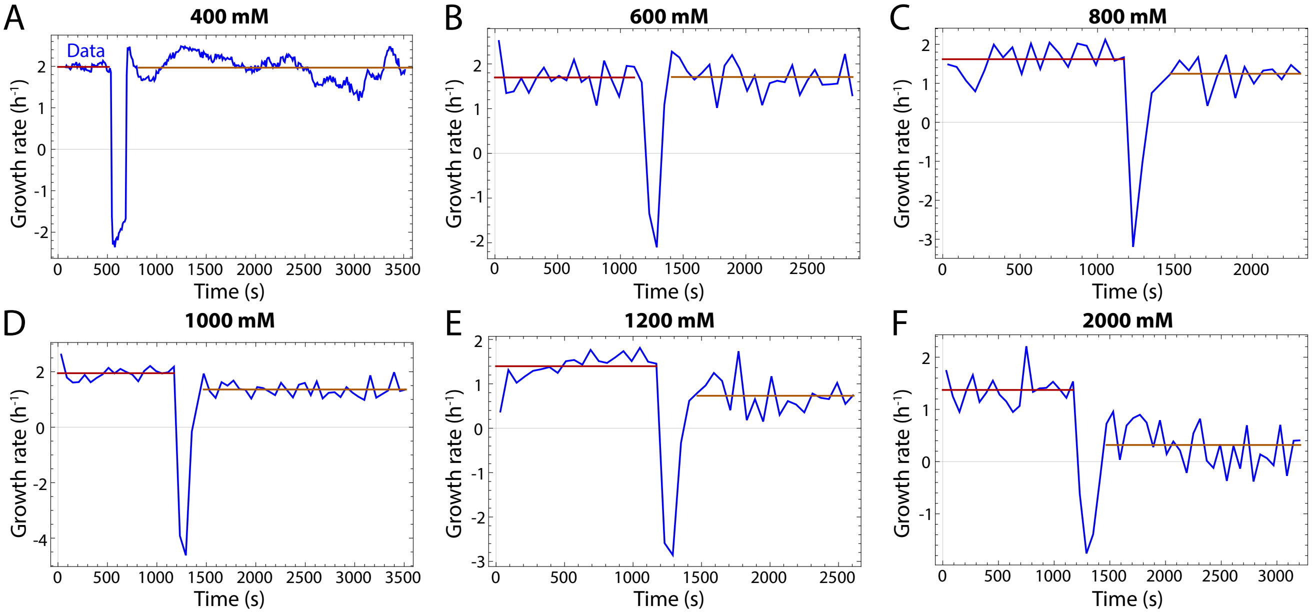


**Fig. S4. Growth rate after an osmotic shock recovers for low shock magnitudes but decreases for large shocks.** The initial growth rate $\lambda_0$ was calculated from all data points before the shock (purple horizontal line), and the final growth rate $\lambda_f$ was calculated after a 5-min interval during which growth rate stabilized after the transient elastic response to the shock. Data in Fig. 4B were derived from these measurements.

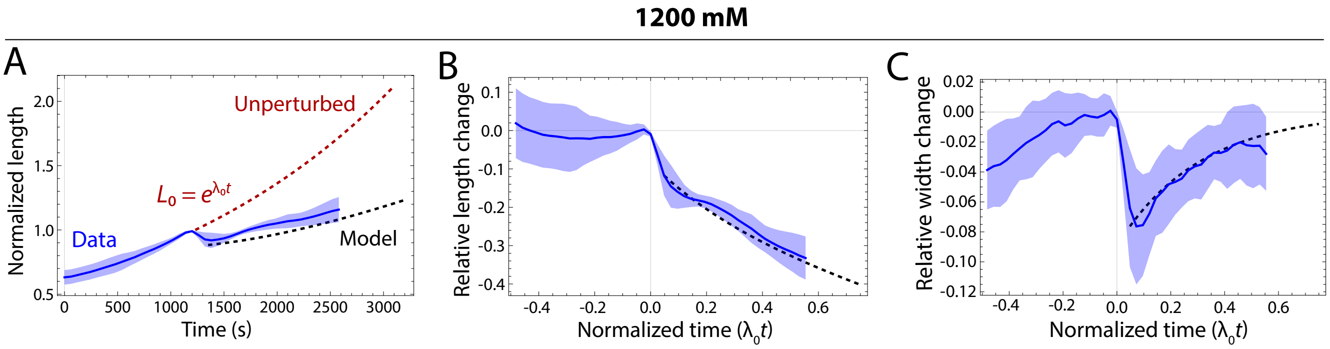


**Fig. S5. Model accurately predicts the response to a 1200-mM hyperosmotic shock.** (A) The predicted dynamics of our model without softening to a 1200-mM hyperosmotic shock (black dashed line) were in close agreement with experimental measurements (blue). The shaded region represents 1 standard deviation. The red curve represents the extrapolated length of an unperturbed cell based on the pre-shock length and growth rate. The parameter estimates $g_1 = 3.2$ and $g_2 = 5.7$ were used. (B) Model predictions of the relative cell length change dynamics were in close agreement with experimental measurements for the data in (A). The shaded region represents 1 standard deviation. (C) Width initially shrank elastically after the hyperosmotic shock due to the decrease in turgor pressure, but then recovered back to its pre-shock value in approximately one cell doubling. The shaded region represents 1 standard deviation. The transient dynamics and recovery are due to the presence of curvature coupling in our model.

where we allowed for the possibility that $u(t)$ is changing with time, for example, due to softening and we defined the parameter combinations

$$g_1 = \frac{1}{4}(\alpha_1 - \alpha_2 - 4), \qquad g_2 = \frac{\alpha_2}{4}, \qquad g_3 = \frac{\beta_1 + 2\beta_2}{2} \tag{S.15}$$

When $g_1 > 0$ and under normal conditions ($u(t) = 1$), the state with constant width $W_0$ and elongation rate $\lambda_0$ is stable: An initial perturbation with $\delta W(0) \neq 0$ and $\delta L(0) \neq 0$ will always evolve towards $\delta W_0(t) = 0$ (or $W(t \to \infty) = W_0$) and $\delta L(t) =$ constant (corresponding to elongation at rate $\lambda_0$).

For a hyperosmotic shock where pressure is reduced to a constant value $p < 1$, and in the absence of softening, we have $u(t) = p$. The length and width change elastically (at fast time scales) in response to this change in pressure, with $\delta L(0) = \delta W(0) = -(1-p)\varepsilon_0$. This case corresponds to equations (6-7) in the Methods.

For constant $u(t) = p$, the solutions of the above equations do not approach $\delta L(t) =$ constant, which continues to decrease due to the term proportional to $(p-1)$. We can determine how the growth rate changes by noting that

$$\lambda(t) \equiv \frac{1}{L(t)}\frac{d}{dt}L(t) = \lambda_0 + \frac{d}{dt}\delta L(t) + O(\delta L)^2 \implies \delta\lambda(t) \equiv \frac{\lambda(t) - \lambda_0}{\lambda_0} \approx \frac{d}{dt}\frac{\delta L(t)}{\lambda_0}. \tag{S.16}$$

In other words, the right-hand side of Eq. (S.14) gives the relative rate change. At long times after a perturbation is applied the width recovers, $\delta W(t \to \infty) = 0$. With $u(t) = p$, we will have $\delta\lambda(t \to \infty) = (p-1)(2+g_3)\varepsilon_0 = (2+g_3)\delta L(0)$. The term $2\delta L(0)$ is simple to interpret geometrically; it represents a constant growth rate per unit area. Therefore, when $g_3 = 0$, changing pressure only affects the growth rate indirectly by changing the area (and curvature).

As described in the main text, softening modifies this behavior so that growth rate recovers its unperturbed value over time (Methods, Eqs. 10-11). Our combined model with stored growth only when $p > 0$ quantitatively explains the behavior of cell after a wide range of shock magnitudes (Figs. 2 - 4 in the main text and S2 - S5).

**Microscopic toy model of the stored growth behavior.** To illustrate stored-growth behavior, we consider a simple model that shows the effects of mechanical tagging of newly inserted material that have distinct mechanical properties and hence can be sensed based on local strains (Fig. 5 in the main text). We assume that at $t = 0$, the cell is a chain of $n_0$ springs with rest length $l_0$ and spring constant $k_0$. The chain thus has an effective Young's modulus $Y_0 = l_0 k_0$. We assume that new springs are always inserted at the same rate $\lambda_0$ so that the total number of springs is given by $n(t) = n_0 \exp(\lambda_0 t)$. However, at $t > 0$, the added springs have different rest length $l_0$ and spring constant $k_1$. Even though the insertion rate is constant before and after the change, the elongation rate will be less than the unperturbed value $\lambda_0$ after the change and will only gradually approach this value over time. This can be seen explicitly through the relation

$$\lambda(t) = \frac{1}{l(t)}\frac{dl(t)}{dt} = \frac{1}{1+e^{-\lambda_0 t}(l_0 - l_1)/l_1}\lambda_0 \qquad \text{[S.17]}$$

where $l(t) = l_1(n(t) - n_0) + l_0 n_0$. Thus, the growth rate approaches $\lambda_0$ from below as the exponential term in the denominator goes to zero.

The microscopic analog of the stored growth condition is

$$\lambda_0\left(1+\frac{F}{Y_0}\right) = \lambda_1\left(1+\frac{F}{Y_1}\right). \qquad \text{[S.18]}$$

This condition ensures that when the applied force F is restored to its initial value, the length of both types of springs are the same. Therefore, this simple toy model illustrates how stored growth behavior can be realized. To calculate the effective Young's modulus of a chain with two types of springs with fractions $n_0/n$ and $n_1/n$, we consider the outcome of stretching the ends with a force F.

Each spring of type i (i=0,1) will stretch by $F/k_i$ to balance the force. The effective Young's modulus is thus

$$Y(t) = Y_0\frac{1-\alpha(l_0-l_1)/l_0}{1+\alpha(k_0-k_1)/k_1}, \qquad \alpha = \frac{n_1(t)}{n(t)} = \frac{l_1(t) - l_0(0)}{l_0(t)}\frac{l_0}{\lambda_1} \qquad \text{[S.19]}$$

Therefore, a measurement of $Y(t)$, also gives a measurement of the deviation from the reference length. This phenomenon is analogous to fluorescently tagging material that is inserted into the chain; by measuring intensity in a particular region on the chain, a change in length could thereby be inferred.
